# LSD1 controls a nuclear checkpoint in Wnt/β-Catenin signaling to regulate muscle stem cell self-renewal

**DOI:** 10.1101/2022.06.10.495614

**Authors:** Sandrine Mouradian, Delia Cicciarello, Nicolas Lacoste, Valérie Risson, Francesca Berretta, Fabien Le Grand, Nicolas Rose, Thomas Simonet, Laurent Schaeffer, Isabella Scionti

**Author notes:** These authors contributed equally (Sandrine Mouradian, Delia Cicciarello, Nicolas Lacoste).

## Abstract

The Wnt/β-Catenin pathway plays a key role in cell fate determination during development and in adult tissue regeneration by stem cells. These processes involve profound gene expression and epigenome remodeling and linking Wnt/β-Catenin signaling to chromatin modifications has been a challenge over the past decades. Functional studies of the histone demethylase LSD1/KDM1A converge to indicate that this epigenetic regulator is a key regulator of cell fate, although the extracellular cues controlling LSD1 action remain largely unknown. Here we show that β-Catenin is a substrate of LSD1. Demethylation by LSD1 prevents β-Catenin degradation thereby maintaining its nuclear levels. Consistently, in absence of LSD1, β-Catenin transcriptional activity is reduced in both MuSCs and ESCs. Moreover, inactivation of LSD1 in mouse muscle stem cells and embryonic stem cells shows that LSD1 promotes mitotic spindle orientation via β-Catenin protein stabilization. Altogether, by inscribing LSD1 and β-Catenin in the same molecular cascade linking extracellular factors to gene expression, our results provide a mechanistic explanation to the similarity of action of canonical Wnt/β-Catenin signaling and LSD1 on stem cell fate.

## Introduction

Resident quiescent muscle stem cells (MuSCs) confer skeletal muscle unique regenerative capacities and play a central role in skeletal muscle plasticity. While quiescent under resting conditions, upon muscle injury MuSCs activate a specific cascade of transcription factors, leave the quiescent state, expand and further differentiate into myocytes, before maturing into myofibers. Notably, a subset of activated MuSCs resists the differentiation process and returns in quiescence to replenish the pool of MuSCs (1–4). The control of the balance between MuSCs commitment and self-renewal is crucial for acute response to muscle injury and to maintain muscle homeostasis overtime.

In order to maintain muscle homeostasis MuSCs, like the majority of stem cells, undergo different modes of cell division: proliferating/self-renewing symmetric division (PSCD), which is the most frequent and gives rise to two stem cells, differentiation symmetric division (DSCD) that gives rise to two committed cells or alternatively an asymmetric division (ACD) that gives rise to a stem cell and a committed cell (5). The choice of ACD over PSCD or DSCD can be controlled by both intrinsic and extrinsic factors. Intrinsic factors generally involve a polarized distribution of cell-fate determinants, such as Par complex, which induce the activation of commitment signals in only one daughter cell (6–8). Consistent with that, perturbation of Par complex in MuSCs results in loss of ACD and reduced regenerative capacity (6, 9). Although these cell-intrinsic mechanisms might be sufficient to achieve asymmetric division, the overall cell polarity is also influenced by external cues, which are generally provided by the environment. Among the extrinsic signals, the secreted Wnt proteins are essential regulator of stem cell division choice. Depending on the Wnt-protein ligand bound to its specific frizzled receptor, three different Wnt signaling pathways have been characterized: the Wnt/β-catenin-, the Wnt/PCP– and the Wnt/Ca^2+^-signaling pathways. However, while the Wnt/Ca^2+^-signaling pathway has never been implicated in the orientation of cell division, the contribution of Wnt/β-catenin– and Wnt/PCP-signaling pathways in favoring ACD has been largely documented in different organisms (8, 10–14). In particular, it has been demonstrated that in mouse embryonic stem cells (ESCs) Wnt3A acts as a positional cue to orient asymmetric cell division (ACD) at single cell level. Such Wnt3A ability rely on the presence of both the Wnt receptors and β-catenin (15). Indeed knocking them out impairs mitotic spindle orientation resulting in equal distribution of cell fate determinants in the daughter cells (14, 15).

The function of the canonical Wnt/β-catenin signaling in regulating stem cells homeostasis has been extensively studied over the past decades. In the absence of Wnt ligands, β-catenin is phosphorylated in the cytosol by the destruction complex (mainly composed by Axin, Adenomatous Polyposis Coli and the Glycogen Synthase Kinase 3 β), which triggers its ubiquitination and rapid degradation by the proteasome. Upon binding of Wnt ligands, such as Wnt3A, to their receptor, the destruction complex is inhibited and β-catenin shuttles into the nucleus and regulates the expression of its-responsive genes. β-catenin protein itself is regulated by multiple post-translational modifications (PTMs), such as phosphorylation, ubiquitinilation or acetylation. Some PTMs enhance and others inhibit β-catenin functions, depending on the nature and the localization of the PTMs (16). In the last decade, β-catenin protein methylation has emerged to be crucial for its transcriptional activity and protein stability both in physiological and pathological conditions (17–21). To date, two methyltransferases have been described to directly modify β-catenin: EZH2 and SET7/9. In particular, while EZH2-mediated tri-methylation of lysine (Lys) 49 represses β-catenin transcriptional activity (18, 19), the SET7/9-mediated mono-methylation of Lys 180 enhances its degradation (20).

Although β-catenin loss or gain of function studies have described severe muscle developmental and regenerative defects, which is its mechanism of action in muscle remains uncharacterized (17, 22).

Lysine specific demethylase 1 (LSD1/KDM1A) is a monoamine oxidase that can de-methylate mono– and di-methylated lysine 4 and 9 residues of the N-terminal of histone H3 (H3K4Me1, H3K4Me2 and H3K9Me1, H3K9Me2) (23, 24). LSD1 can also demethylate non-histone proteins influencing their transcriptional activity (P53) (25) or protein stability (HIF-1α and DNMT1) (26, 27). Altogether, these studies indicate that LSD1 can regulate gene transcription by acting both on histones, on transcription factors and DNA methylation. We have previously shown that LSD1 enzymatic activity is crucial for the timely expression of *MyoD* during the commitment of muscle progenitors, via the transcriptional activation of the Core Enhancer Region (CER) of the *MyoD* gene (28). Consistently, LSD1 ablation delays *MyoD* expression during embryonic myogenesis.

LSD1 and Wnt/β-catenin are both key players in the regulation of progenitor cell stemness and engagement of progenitor cells into differentiation (29, 30). However, functional and molecular link between them were not identified yet.

Here we demonstrate that β-catenin is a non-histone target of LSD1, by showing that once in the nucleus β-catenin needs to be demethylated by LSD1 to prevent its degradation and activate the transcription of its target genes. Moreover, we demonstrate that LSD1 and β-catenin work together to regulate the mitotic spindle orientation, both in MuSCs and ESCs.

## Materials and Methods

### Animals

All procedure on animals were performed in accordance with European regulations on animal experimentation and were approved by the local animal ethics committee (CECCAPP, University of Lyon) under the reference Apafis#16930. Mice were bred and housed in AniRA-PBES animal facility. They were maintained in a temperature– and humidity-controlled facility with a 12h light/dark cycle, free access to water and standard rodent show.

Nine-week-old C57BL/6J mice were purchased from Charles River laboratories and intraperitoneal injected with OG-L002 20mg/kg or NaCl 0,9% for 7 days.

The Lsd1^tm1Schüle^ (31) and Pax7-CreERT2 (32) were previously described. Mice were genotyped with conventional PCR using standard conditions.

Allelic recombination under the Pax7-CreERT2 allele was induced by intraperitoneal injections of 2mg of Tamoxifen (TAM) for 5 days.

Skeletal muscle injury was induced by an injection of 50 μl of cardiotoxin (10μM) into hindlimb TA muscle using 30G syringes under anesthesia induced by intraperitoneal injection with Ketamine (100mg/kg) and Xylazine (10mg/kg) in sterile saline solution. EdU solution (200μg) was injected intraperitoneally 12 hours before euthanasia.

### Cell cultures

C2C12 mouse myoblasts were maintained as myoblasts in growth medium (GM): Dulbecco’s modified Eagle’s medium supplemented with 15% fetal calf serum and antibiotics. Adult primary MuSCs were maintained on Matrigel-coated dishes in GM: Dulbecco’s modified Eagle’s medium F12 supplemented with 20% horse serum, 5 ng/mL fibroblast growth factor (FGF), and antibiotics. C2C12 cells and adult primary MuSCs were differentiated into myotubes by replacing GM with a myogenic differentiation medium containing 2% horse serum with antibiotics (MDM). HEK 293T cells were cultured using standard methods (ATCC).

### Mouse embryonic stem cells (ESCs)

To generate LSD1 knock-out (KO) ESCs (referred to LSD1 KO), the Gene Knockout Kit v2 (Synthego Corp), which comprises three strategically designed modified sgRNAs inducing fragment deletion (sgRNAs LSD1 sequences: UCCAUGGGCGUCGCGGAGCC, CCGAGCCGCCGGGGUCUGCU, and CGAGCGCACUCCCCGAAAGA) has been used. The ribonucleoprotein (RNP) assembly was conducted immediately before electroporation of ESC. In brief, 102 pmol of Streptococcus Pyogenes protein (Alt-R® S.p. Cas9 Nuclease V3, cat. num. 1081058, IDT) was mixed with 133 pmol of synthetic sgRNAs (Gene Knockout Kit v2, Synthego Corp) and incubated at room temperature for 20 minutes. 1 × 10^6^ ESCs were suspended in 100 μL of OptiMEM buffer and added to the RNP solution, along with a final concentration of 1.8 μM of Electroporation Enhancer (IDT). The mixture was gently combined, and electroporation was conducted using a NEPA21 electroporator (Nepa Gene Co., Ltd.) with 2mm gap cuvettes in the 100 µL format. NEPA21 generates a poring pulse (Pp) and a transfer pulse (Tp). The electroporation parameters were set as follows: 2 Pp (125 V, 5 ms pulse duration, 50 ms pulse interval, 10% decay (+ pulse orientation)) and 5 Tp (20 V, 50 ms pulse duration, 50 ms pulse interval, 40% decay (± pulse orientation)). After electroporation, cells were seeded into 100 mm dishes. Monoclonal cell lines were produced through limiting dilution cloning into two gelatin-coated p96 plates. LSD1 KO clones were screened using Western blot analysis with antibody against LSD1.

Wild-type mouse ESCs (with a 129 background) heterozygous β-catenin deficient (β-cat fl/-) and LSD1 KO ESCs were cultured on 0.2% gelatin-coated plates at 37°C in 5% CO2 in 2i/LIF medium composed of: Advanced DMEM/F12/Glutamax, 10% ESC-qualified FBS, 1% penicillin-streptomycin, 50 μM β-mercaptoethanol and 1000 U/mL recombinant Leukaemia Inhibitory Factor. This medium was supplemented with 2i: 1 μM MEK inhibitor PD0325901 and 3 μM GSK3 inhibitor CHIR99021. Cell confluency was maintained below 80% and regularly tested for the absence of mycoplasma.

To induce β-catenin knockout in βfl/-cells, 0.1 mg/mL (or 258 nM) 4-hydroxy-tamoxifen (4-OHT) was added to the media every 24 hours for 72 hours. Cells treated with 4-OHT for 3 days were used in the experiments (referred to as β-Cat KO), while cells treated with DMSO served as controls.

### MuSCs isolation

Adult primary MuSCs were isolated from skeletal muscle tissue as previously described (33). Digested tissue was stained using antibodies: 0.2μg PE-conjugated rat anti-mouse CD31, 0.2μg PE-conjugated rat anti-mouse CD45, 0.2μg PE-conjugated rat anti-mouse Sca-1, 5μg 647-conjugated rat anti-mouse integrin alpha7 and 2.5μg rat anti-mouse CD34. Cells were incubated with primary antibodies for 40 minutes on ice. Adult primary MuSCs were FACS isolated into MuSC medium based on cell surface antigen markers: CD31-/CD45-/Sca1-/integrin-ɑ7+/CD34+ using a FACsAria III.

### Myofiber culture

We performed myofiber culture as described earlier (34). Briefly, we carefully dissected Extensor Digitorum Longus (EDL) muscles and incubated them in DMEM (Gibco) containing 600U/ml of collagenase B for 75 min. The myofibers were dissociated by gentle trituration with a glass pipette. Myofibers were cultured for 42 h in DMEM containing 20% FBS and 5 ng/ml bFGF.

### Cell treatments

Inhibition of the proteasome was carried out by treating HEK 293T cells with MG132 at the indicated concentration and incubated for 6h at 37°C before the collection of cells. In protein degradation assays, protein synthesis was inhibited by the addition of cycloheximide (CHX) to C2C12 stable clones after 72h in MDM at a final concentration of 50 mg/ml in a time course before harvest, as indicated in the text. Canonical Wnt pathway activation was achieved by treating C2C12 stable clones and primary adult MuSCs with Wnt3A at 50 ng/ml or LiCl at 25mM for 6 h before the luciferase assay, proximity ligation assay, or RNA extraction. Inhibition of LSD1 enzymatic activity was carried out by treating primary adult MuSCs with Pargyline at 1mM and OG-L002 at 10μM for 72h in MDM. For proliferation assay EdU solution (20mM) has added to adult primary MuSCs for 2 hours before the collection of cells. Cells were analyzed using flow cytometry (FACS Cantoll). For Wnt3A-dependent cell division assay, freshly isolated CTRL SC or LSD1 SCiKO MuSCs were exposed to Wnt3A-coupled to beads and analyzed as previously described (15).

### Cell Transfection

Different plasmids were transfected into C2C12 stable clones, MuSCs or HEK 293T with lipofectamine or JetPRIME following provider’s instruction for luciferase assays, immunoblotting, Co-IP, Demethylase assay or assessment of Wnt3A-cell division. To perform luciferase assays, C2C12 stable clones and primary adult MuSCs were grown in 6-well plates and cells in each well were co-transfected with 100 ng of TopFlash or FopFlash reporter plasmid and 1 ng of pRL-TK Renilla luciferase reporter. For immunoblotting analysis that was used for detecting the proteasome dependent degradation of β-Catenin protein and its mutant, 2 μg of pCMV β-Catenin WT or its mutant pCMV β-Catenin K180R were transfected into HEK 293T cells, which were cultured at about 70-80% confluency in 3.5 cm dishes. For Co-IP experiments, pCMV-LSD1 flag or pCMV β-Catenin or pCMV GFP were transfected into 70-80% confluent HEK 293T cells that were cultured in 15-cm culture dishes. For demethylase assay, 15 cm plates containing 70-80% confluent HEK 293T cells were transfected with pCMV-LSD1 flag or pCMV-LSD1 K661A flag or pCMV-LSD1 K661A W754A Y761A flag or pCMV-GFP (CTRL) after 24h incubation, protein extraction has been performed. For Wnt3A-dependent cell division assay, freshly isolated CTRL SC or LSD1 SCiKO MuSCs were transfected with an shRNA against β-catenin or a *wild-type*-β-catenin fused with mCherry respectively, after 24h, cells were exposed to Wnt3A-coupled to beads and analyzed as previously described (15).

### Luciferase assay

C2C12 stable clones and primary adult MuSCs were transfected as previously described. After 24h, Wnt3A was added and incubated for another 6h at 37°C and luciferase activity was measured using the Dual-Luciferase Assay System (Promega). Each measurement was repeated with at least three independent transfections.

### Immunoprecipitation

Protein complexes were precipitated from nuclear fraction. Cell fractionation was performed as described (35). 500μg of nuclear extracts from C2C12 stable clones, pCMV-LSD1 flag or pCMV-GFP transfected cells were incubated with specific antibodies coupled with beads or Flag beads in IP buffer (20mM HEPES pH7.5, 5mM K acetate, 0.5mM MgCl2, 0.5mM DTT, 150mM NaCl, 0.5% NP40 and Complete protease inhibitor) overnight at 4°C on a wheel. Beads were then washed 3 times in the IP buffer and resuspended in 1X loading sample buffer (IP buffer + 50mM Tris HCl pH 6.8, 10% glycerol, 100mM DTT, 2% SDS and bromophenol blue). Proteins were then loaded on a 4-15% SDS-PAGE transferred on a nitrocellulose membrane and blotted against the indicated antibodies.

### Immunoblotting

Cell fractionation was performed as described (35) and quantified using the DC protein assay (Bio-Rad). Nuclear extracts were separated by electrophoresis on 10% SDS-PAGE and transferred onto polyvinylidene fluoride (PVDF) Immobilon-P membranes. Immunoblots were revealed with enhanced chemiluminescence (ECL) PLUS reagent (Cytiva) according to the manufacturer’s instructions.

### Immunofluorescence

For plated MuSCs, 4 well Permanox chamber slides (Nunc Lab-Tek) were used and cells fixed with 4% (v/v) paraformaldehyde (PFA) for 10 minutes. Following fixation, material was permeabilized with 0.5% (v/v) Triton X-100 solution for 5 minutes and then blocked with 1% BSA, 0.2% Triton X-100 and 5% (v/v) serum for 60 minutes to reduce nonspecific antibody binding. Cells were then incubated with the following antibodies overnight at 4^°^C, 1:50 mouse anti-PAX7 (DSHB, clone PAX7), 1:50 mouse anti-Myogenin (DSHB, clone FD5), 1:50 mouse anti-MyHC (DSHB, clone A4.1025). Species-specific fluorochrome-conjugated secondary antibodies were then applied for 1 h at room temperature, before being mounted with 100 ng/ml of DAPI.

### Chromatin immunoprecipitation (ChIP)

ChIP experiments were carried out essentially as previously described (28).

### *In vitro* Demethylation assay

HEK 293T cells were transfected with pCMV-LSD1 flag or pCMV-LSD1 K661A flag or pCMV-LSD1 K661A W754A Y761A flag or pCMV-GFP (CTRL). After 24h incubation, 4 mg of nuclear extracts were mixed with 40μl of Flag beads in IP buffer (20mM Tris Hcl pH8, 300mM NaCl, 0.5% NP40, 5% glycerol and Complete protease inhibitor during 3h at 4°C on a wheel. Beads were then washed 3 times in the IP buffer and eluted 2 times 90 min at 4°C on a wheel with 40μl of elution buffer (IP buffer + 150μg flag peptide). 15μl of each purification was used for the demethylase assay using 50μM (final concentration) of β-CAT WT peptide or β-CAT K180me peptide in 1X demethylase buffer (50mM Tris HCl pH8, 50nM FAD, 50mM NaCl, 5% glycerol, 20μg/ml BSA). 0.8μg of recombinant LSD1 (Sigma-Aldrich SRP0122) was used as positive control. After 1h at 37°C the reaction was stopped by adding sample loading buffer (final concentration 50mM Tris HCl pH 6.8, 10% glycerol, 100mM DTT, 2% SDS and bromophenol blue) and half of the reaction was loaded on a 4-20% precast gel. After transfer on a 0.2μm PVDF membrane proteins are detected with the indicated antibodies and with streptavidin coupled with HRP for the biotinylated peptides as loading control.

### Real-Time qPCR

Total RNA was isolated from cultured cells at 72h in MDM grown in 100-mm dishes using Tri Reagent (Sigma). RNA was analyzed by real-time PCR using the QuantiFast SYBR Green PCR Kit. Relative gene expression was determined using the DCt method.

### Proximity ligation Assay (PLA)

Duolink® *in situ* PLA reagents were used to detect the interaction between β-catenin and BCL9 and the manufacturer’s protocol was followed. Briefly, MuSCs, cultured on chamber slides to 40-50% confluence, were treated with Wnt3a (50ng/mL for 6 hours). Cells were treated with cytoskeleton buffer (PIPES 10mM, NaCl 100mM, Sucrose 300mM, MgCl_2_ 3mM, EgTA 1mM and TrotonX100 0.5%), washed with PBS and then treated with cyto-stripping buffer (Tris-HCl 10mM, NaCl 10mM, MgCl_2_ 3mM, Tween 40 1% and Sodium deoxycholate 0.5%) to remove the cytoplasm. Cells were then fixed, permeabilized and incubated overnight with mouse anti-β-catenin and rabbit anti-BCL9. The following day, cells were incubated with secondary antibodies conjugated to oligonucleotides (PLA probe PLUS and PLA probe MINUS) for 1 hour at 37°C. Afterward, ligation solution containing 2 oligonucleotides (that hybridize to the PLA probes) and Ligase was added for 30 minutes at 37°C. A closed circle is only formed if the 2 PLA probes are in close proximity. Finally, the closed circle was amplified using rolling-circle amplification reaction and the product was hybridized to fluorescently-labeled oligonucleotides. The fluorescent spots generated from positive interactions were quantified using a confocal microscope (Leica TCS SP5).

### Muscle Histology and immunohistochemistry

Mice were euthanized and TA muscles were dissected and attached in Tragacanth gum, frozen in a cold 2-methylbutane bath and cryo-sectioned onto glass slides.

Tissue sections were freshly fixed with PFA 4% for 10 minutes, washed three washes in PBS, permeabilized in methanol for 6 minutes at –20°C and then washed three times in PBS. After 10 minutes incubation in a hot antigen retrieval buffer, sections were washed three times in PBS, 0,1% triton X-100 (PBS-T) and then were saturated 2 hours at room temperature with M.O.M Mouse IgG Blocking reagent. The antigen retrieval buffer contained 10mM sodium citrate acid and 0,05% tween-20 and was adjusted at pH 6,0. Tissue sections were washed once in PBS-T, incubated for 5 minutes in M.O.M diluent prepared according to the manufacturer and stained at 4°C overnight with primary antibodies diluted in M.O.M diluent (1/200 for anti-Ki67, 1/800 for anti-LSD1, 1/200 for anti-laminin, 1/50 for anti-Myogenin, 1/50 for anti-Pax7. Pax7 antibody has been obtained by concentrated 33 times the supernatant of the hybridoma’s culture). After three 10 minutes washes in PBS-T, sections were incubated for 1 hour at room temperature with secondary antibody and DAPI diluted in M.O.M diluent. After three washes, sections were mounted with Fluoromount-G. Fluorescent images were acquired on a Zeiss Z1 Axioscan.

Hematoxylin and Eosin (H&E) and Oil Red O (ORO) staining were performed following standard methods.

## Results

### LSD1 inactivation increases MuSCs pool upon muscle injury

To investigate the role of LSD1 in adult MuSCs, LSD1 was specifically inactivated in MuSCs using LSD1 conditional Knock-Out mice expressing a tamoxifen-inducible CRE-recombinase under the control of the *Pax7* promoter (hereafter named LSD1 SCiKO mice). 10 weeks old mice were treated with tamoxifen (TAM), and MuSCs were FACS-isolated 1 week later. MuSCs isolated from Pax7-CreERT2 mice treated with TAM (hereafter named CTRL SC) were used as a control (Supplementary Figure S1A). A high degree of recombination was evident in FACS-purified MuSCs from LSD1 SCiKO mice based on the amount of LSD1 protein level (Supplementary Figure S1B). As previously reported (28), the ablation of LSD1 did not impact the percentage of MuSCs in S-phase *in vitro* (Supplementary Figure S1C). After 2 days in myogenic differentiation medium (MDM) LSD1 SCiKO MuSCs displayed impaired myogenesis as evidenced by a strong reduction in the proportion of Myogenin positive (+) cells compared to CTRL SC MuSCs (Supplementary Figure S1D). Conversely, the proportion of PAX7+ cells was significantly increased in LSD1 SCiKO MuSCs compared to CTRL SC MuSCs (Figure 1A). This increase was caused by the loss of LSD1 enzymatic activity since the LSD1 pharmacological inhibitors Pargyline and OG-L002 produced the same effect on control MuSCs (Figure 1B). Altogether, these results suggested that the loss of LSD1 delayed MuSCs commitment as we previously showed for myogenic progenitors during development (28). Reduced myogenesis could not be attributed to activation of the adipogenic program, since LSD1 SCiKO MuSCs did not produce lipid-containing mononucleated cells in MDM (Figure 1C).

**Figure 1.**
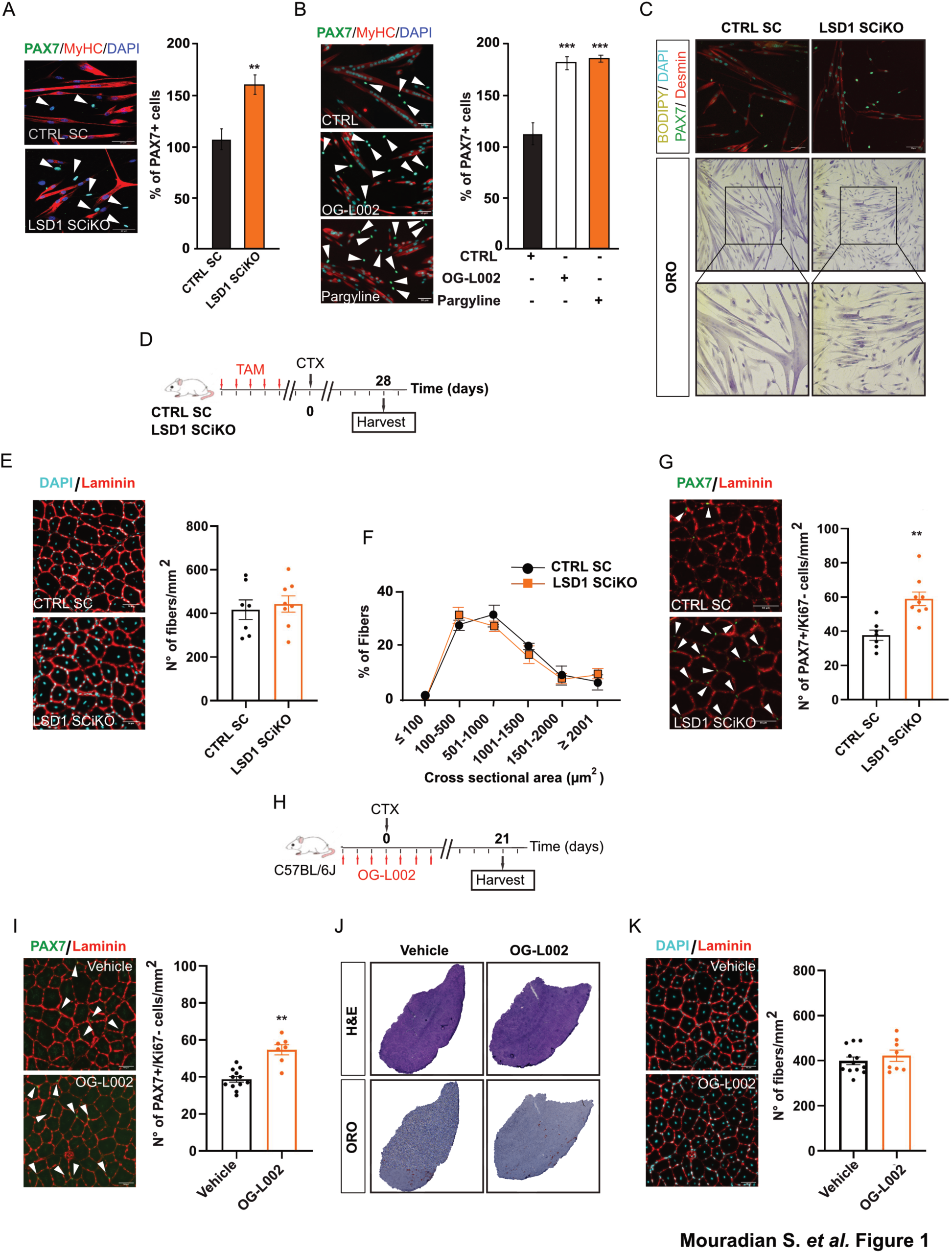
LSD1 regulates MuSCs self-renewal potential. (**A**) PAX7 and MyHC staining and percentage of PAX7+/Ki67-cells after 48 h under myogenic differentiation conditions of CTRL SC and LSD1 SCiKO MuSCs. (**B**) PAX7 and MyHC staining and percentage of PAX7+/Ki67-cells after 48 h under myogenic differentiation conditions of MuSCs treated with LSD1 inhibitors (OG-L002 and Pargyline). (**C**) Bodipy, PAX7, desmin and Oil Red O staining of cells after 48 h under myogenic differentiation conditions of CTRL SC and LSD1 SCiKO MuSCs. (**D**) CTX experimental setup. (**E**) Anti-PAX7 and anti-Laminin staining on cryosections of regenerated TA muscles in CTRL SC and LSD1 SCiKO mice at 28 dpi. Quantification of the number of sublaminar PAX7+/Ki67-cells per mm^2^. (**F**) Anti-Laminin staining on cryosections of regenerated TA muscles in CTRL SC and LSD1 SCiKO mice at 28 dpi. Quantification of the number of myofibers per mm^2^. (**G**) CSA distribution of muscle fibers in CTRL SC and LSD1 SCiKO mice TA cryosections at 28 dpi. (**H**) CTX experimental setup. (**I**) Anti-PAX7 and anti-Laminin staining on cryosections of regenerated TA muscles in Vehicle and OG-L002 treated mice at 21 dpi. Quantification of the number of sublaminar PAX7+/Ki67-cells per mm^2^. (**J**) H&E and Oil Red O staining on cryosections of regenerated TA muscles in Vehicle and OG-L002 treated mice at 21 dpi. (**K**) Anti-Laminin staining on cryosections of regenerated TA muscles in Vehicle and OG-L002 treated mice at 21 dpi. Quantification of the number of myofibers per mm^2^. Scale bars, 50 µm. n = 3 mice/genotype. n = 3 primary MuSC cultures/genotype. n = 3 primary MuSC cultures/treatments. Values are mean or percentage mean ± SEM. **p < 0.01 (Mann-Whitney-Wilcoxon Test) ***p < 0.001 (Bonferroni test after one way-ANOVA).

We next examined the effect of LSD1 inactivation on MuSCs *in vivo*. *Lsd1* gene inactivation did not affect the number of quiescent MuSC in healthy muscles, as shown by PAX7 immunofluorescence on *Tibialis Anterior* (TA) muscle cryosections (Supplementary Figure S2A, B). Of note, 7 and 56 days after the last TAM injection, the number of fibers per mm^2^ and the muscle histological features (Supplementary Figure S2C, D) were also similar in LSD1 SCiKO and CTRL SC uninjured TA muscles. To investigate LSD1 role during muscle regeneration, TA muscles of LSD1 SCiKO and CTRL SC were injured by single cardiotoxin (CTX) injection. Seven days after CTX injection (Supplementary Figure S2E), LSD1 SCiKO TA muscles displayed more smaller regenerating fibers than CTRL SC TA muscles (Supplementary Figure S2F, G). Surprisingly, 28 days post injury (dpi), when muscles were completely regenerated (Figure 1D), LSD1 SCiKO TA muscles showed similar fibers size and density (number of fibers per mm^2^) than CTRL SC TA muscles (Figure 1E, F). Consistently with the *in vitro* results, we did not observe an increase in adipocyte-like cells (Supplementary Figure S2H). These results suggest that in absence of LSD1, while muscle regeneration is initially impaired, it ultimately occurs. Consistently, we quantified the number of MYOG+ cells at 7dpi and we observed a significant increase of MYOG+ myocytes in LSD1 SCiKO TA muscles compared to CTRL SC, indicating that MuSCs differentiation was delayed (Supplementary Figure S2I). Remarkably, at 28 dpi, when MuSCs had returned into quiescence, the number of PAX7+ cells per mm^2^ was significantly increased by 50% in LSD1 SCiKO muscles compared to CTRL SC (Figure 1G). Staining for PAX7 and the proliferation marker Ki67 showed that in both conditions more than 95% of the PAX7+ cells were negative for Ki67, indicating that they were not proliferating (Supplementary Figure S2J). Altogether, these results indicate that in absence of LSD1 the total number of MuSCs increases during the regeneration process.

To investigate whether pharmacological inhibition of LSD1 produced similar effects, *wild-type* (WT) mice were treated with the pharmacological inhibitor of LSD1 (OG-L002 (36)) and TA muscles were injured by single CTX injection (Figure 1H). Twenty-one dpi, the number of quiescent MuSCs (PAX7+/Ki67-) in TA muscles was significantly higher in mice treated with OG-L002, compared to vehicle (Figure 1I). The histological analysis and the number of fibers per mm^2^ showed no significant differences between treated and untreated mice (Figure 1J, K). These results support the conclusion that the effect of LSD1 inactivation on MuSCs function is due to the loss of LSD1 enzymatic activity.

### MuSCs lacking LSD1 maintain their regenerative and self-renewal potential after repeated injury

Since LSD1 ablation or enzymatic inhibition increased the pool of MuSCs, we hypothesized that it could either be beneficial or detrimental for further regeneration events. Three consecutive muscle regenerations were therefore performed by three successive CTX injections 28 days apart from each other in LSD1 SCiKO and CTRL SC TA muscles. Injured muscles were analyzed 7 and 28 days after the third CTX injection (dpiIII, Figure 2A). Surprisingly, fiber number per mm^2^ were not reduced anymore at 7 dpiIII in LSD1 SCiKO conversely to what was observed after a single round of regeneration (Figure 2B and Supplementary Figure S2F). In addition, regeneration efficiency at 28 dpiIII was similar to what was observed after a single CTX injection in CTRL SC: the distribution of regenerated fibers CSA, the number of fibers for mm^2^ as well as the histological analysis was comparable between CTRL SC and LSD1 SCiKO mice (Figure 2B-D) but a 40% increase in the number of self-renewed MuSCs (PAX7+/Ki67-)/mm^2^ in LSD1 SCiKO mice compared to CTRL SC was still present (Figure 2E). These results indicated that the loss of LSD1 in MuSCs increased the pool of PAX7+ while preserving the regenerative potential of muscles even after repeated injuries.

**Figure 2.**
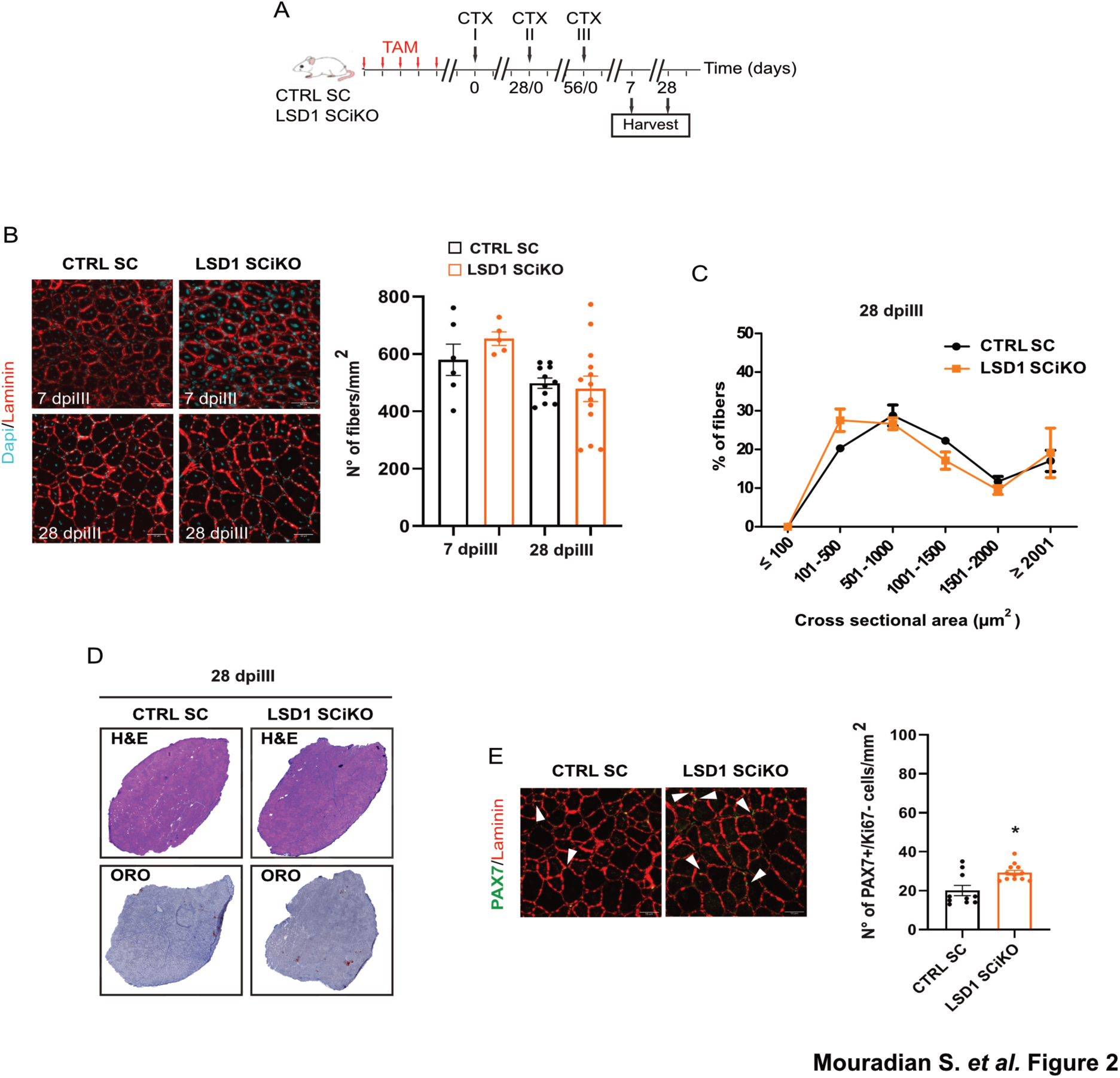
LSD1 SCiKO MuSCs maintain their regenerative potential after repeated injuries. (**A**) Repeated CTX experimental setup. (**B**) Laminin staining on cryosections of regenerated TA muscles in CTRL SC and LSD1 SCiKO mice at 7 and 28 dpiIII. Quantification of the number of myofibers per mm^2^. (**C**) CSA distribution of muscle fibers in CTRL SC and LSD1 SCiKO mice TA cryosections at 28 dpiIII. (**D**) H&E and Oil Red O staining on cryosections of regenerated TA muscles in CTRL SC and LSD1 SCiKO mice at 28 dpiIII. (**E**) Anti-PAX7 and anti-Laminin staining on cryosections of regenerated TA muscles in CTRL SC and LSD1 SCiKO mice at 28 dpiIII. Quantification of the number of sublaminar PAX7+/Ki67-cells per mm^2^. Scale bars, 50 µm. n = 5 mice/genotype. Values are mean ± SEM. *p < 0.05 (Mann-Whitney-Wilcoxon Test).

### LSD1 is critical for asymmetric cell division

To better understand the mechanism underlying this phenotype, we investigated at which specific stage LSD1 inactivation affected MuSCs amplification *in vivo*. After muscle injury, MuSCs undergo a first round of division between 28 and 40 hours (h) (37, 38). A pulse of the deoxynucleotide analog EdU labeling was performed 28 h post injury (hpi). MuSCs were isolated 12 h later (40 hpi) and EdU incorporation was analyzed by cytometry (Figure 3A). MuSCs were activated similarly in LSD1 SCiKO and CTRL SC muscles (Figure 3B).

**Figure 3.**
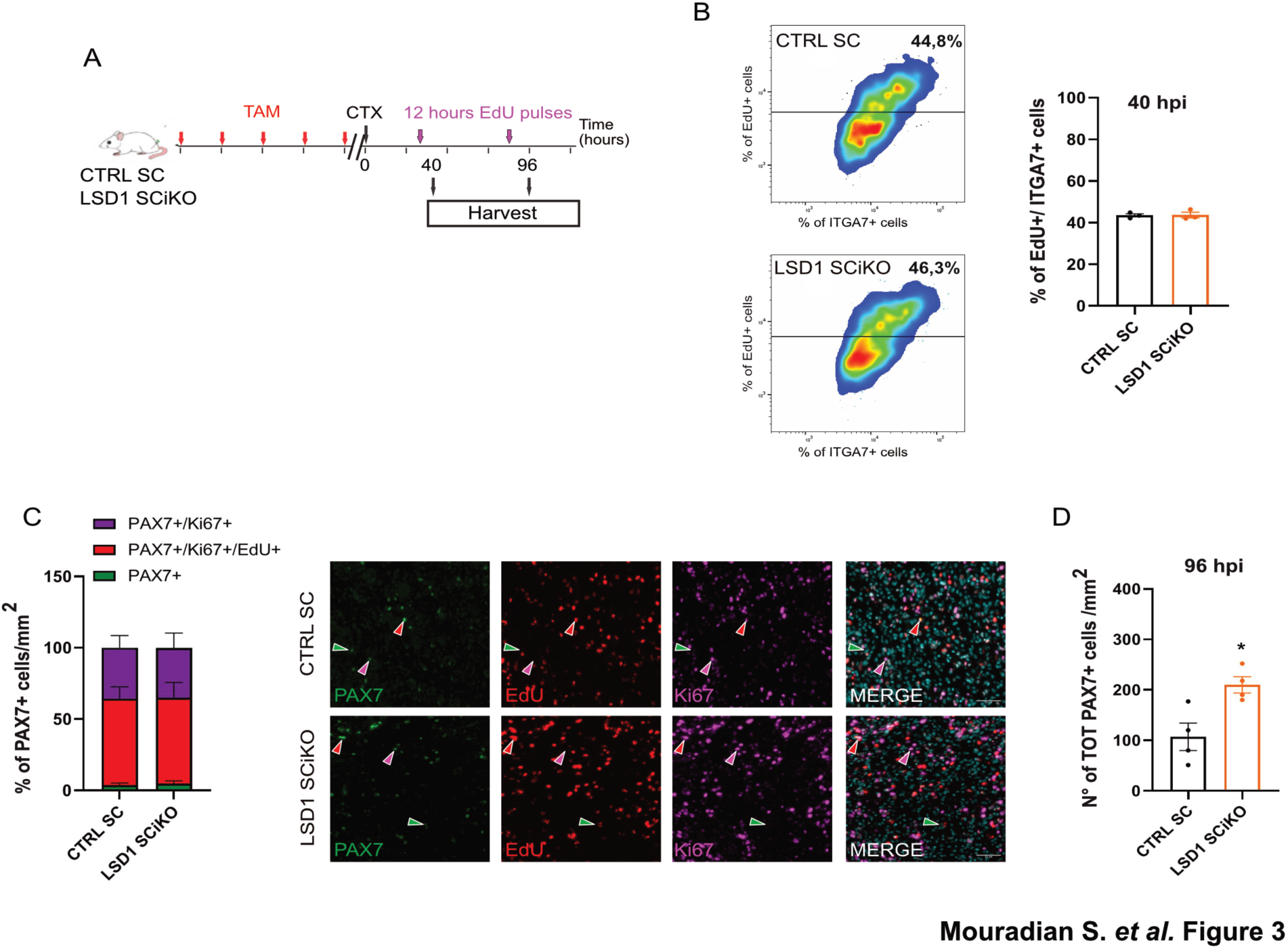
Lack of LSD1 increases MuSC pool upon injury. (**A**) EdU pulse labelling set up. (**B**) FACS-acquired quantification of EdU+ MuSCs (ITGA7+) isolated from muscle 40 h post injury (hpi). (**C**) Percentage of PAX7+/Ki67+ (violet), PAX7+/Ki67+/EdU+ (red) and PAX7+/Ki67-/EdU-(green) MuSCs at 96hpi was quantified per mm^2^. Representative images of PAX7, Ki67 and EdU staining on cryosections of regenerated TA muscles. (**D**) Quantification of the total number of PAX7+ cells at 96 hpi per mm^2^. Scale bars, 50 µm and 10 µm. n = 3 mice/genotype. Values are mean or percentage mean ± SEM. *p < 0.05 (Mann-Whitney-Wilcoxon Test).

After completing the first round of division MuSCs rapidly proliferate to expand their number to allow muscle repair. To measure the rate of MuSC expansion, muscles were analyzed by PAX7, Ki67 immunostaining and EdU incorporation at 96 h post injury (Figure 3A, C). No difference was observed between the proliferation rate of CTRL SC and LSD1 SCiKO MuSCs (Figure 3C). However, the absolute number of PAX7+ cells was already significantly increased in absence of LSD1 (Figure 3D and Supplementary figure S2K). Altogether, our data suggested that LSD1 might be involved in the regulation of the balance between proliferation/self-renewing symmetric (PSCDs) and asymmetric divisions (ACDs). To further investigate this hypothesis, myofibers were isolated from LSD1 SCiKO and CTRL SC EDL muscles and placed in culture for 42 h before immunolabelling stem cells with an anti PAX7 antibody and committing cells with an anti MYF5 antibody, as previously described (39). In the absence of LSD1, ACDs were drastically reduced as evidenced by the percentage of cell doublets with an asymmetric expression of the commitment marker MYF5 that dropped to 2% compared to the 15% observed in CTRL SC (Figure 4A). Conversely, the frequency of PSDC was significantly increased to 98% in LSD1 SCiKO MuSCs (Figure 4A). These data indicated that LSD1 favors asymmetric divisions during the first division of MuSCs.

**Figure 4.**
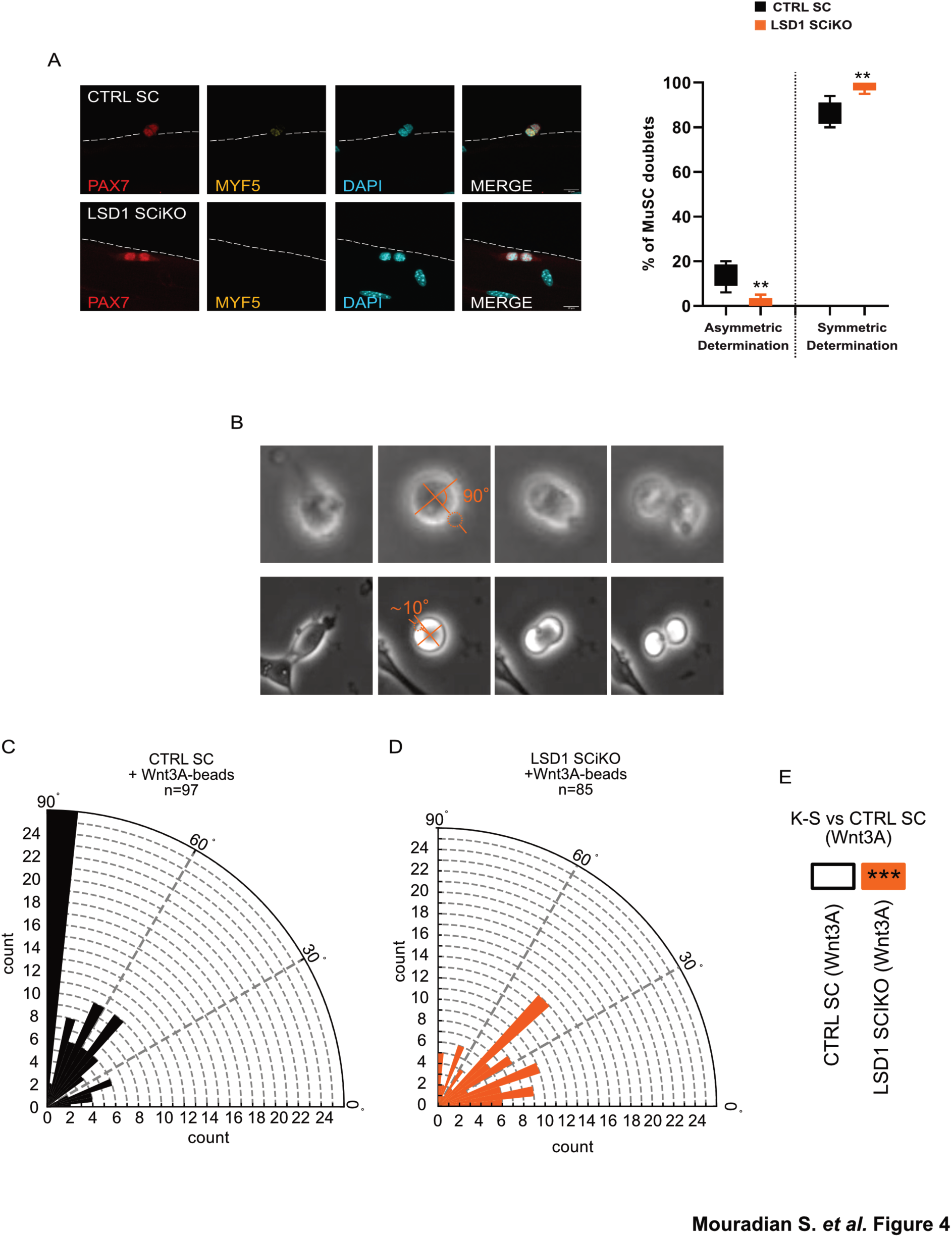
Lack of LSD1 stimulates symmetric division. (**A**) Representative images of MuSC doublets. Anti MYF5 (Yellow) and PAX7 (Red) staining on EDL myofibers isolated from CTRL SC and LSD1 SCiKO mice and cultured for 42 hours. Percentage of doublets of CTRL SC or LSD1 SCiKO MuSCs. Asymmetric determination = PAX7+/MYF5+; Symmetric determination = PAX7+/MYF5-. n = 6 mouse/genotype. Values are percentage mean ± SEM. **p < 0.001 (Mann-Whitney-Wilcoxon Test). (**B**) Representative images of two MuSCs dividing with Wnt3a-beads at angles of ∼10° and 90°. Orthogonal orange lines depict major and minor axis, orange line points to Wnt3a-bead and shows the angle measured. Wnt3a-bead is highlighted with orange dashed circle. (**C-D**) Rose plots illustrating the distribution of mitotic spindle angles in CTRL SC MuSCs (**C**) or LSD1 SCiKO MuSCs (**D**), dividing with Wnt3a-beads. n= number of cells analyzed. (**E**) Statistical comparison of the distribution of mitotic spindle angles in LSD1 SCiKO MuSCs to the one observed in CTRL SC MuSCs. ***p < 0.001 (Kolmogorov-Smirnov test)

To confirm this phenotype, we exposed freshly isolated MuSCs to Wnt3A molecules covalently attached to carboxylic acid-coated beads, and we measured the orientation of the mitotic spindle in relation to the position of the beads, as previously described in ESCs (15, 40) (Figure 4B). CTRL SC MuSCs mitotic spindle was mainly oriented toward the Wnt3A-coupled beads with an angle of 85-90° (Figure 4C). Conversely, LSD1 SCiKO MuSCs displayed an impairment in Wnt3A-dependent mitotic spindle orientation (Figure 4D). The majority of LSD1 SCiKO MuSCs divided with angles lower than 50° compared to CTRL SC MuSCs (Figure 4E). Our results suggest that LSD1 is involved in Wnt3A-dependent mitotic spindle orientation.

### LSD1 is required for transcriptional activation by β-catenin

To determine whether LSD1 function could be linked to Wnt/β-catenin signaling, shLSD1 and control shSCRA myoblasts (28) as well as CTRL SC and LSD1 SCiKO primary MuSCs were transfected with a β-catenin reporter luciferase construct and treated with Wnt3A or LiCl (41) to activate Wnt/β-catenin signaling. In absence of LSD1, myoblasts displayed very low luciferase expression compared to control cells (Supplementary Figure S3A). Next, we evaluated the effect of LiCl on the activation of β-catenin target genes and as expected, LiCl increased the expression of *Fst*, *Porcn* and *Axin2* in shSCRA cells whereas their expression was hardly increased in shLSD1 cells (Supplementary Figure S3B).

To promote myogenic cells differentiation, the Wnt/β-catenin signaling requires the formation of a complex between β-catenin and BCL9 (17, 42). Proximity ligation assay (PLA) was used to evaluate the interaction between β-catenin and BCL9 in CTRL SC and LSD1 SCiKO MuSC treated with Wnt3A. The results indicated that significantly less β-catenin/BCL9 complexes were formed in LSD1 SCiKO than in CTRL SC MuSCs (Figure 5A). Altogether, these results showed that LSD1 is required for the activation of gene expression by β-catenin in myogenic cells. Importantly, LiCl treated shLSD1 myocytes expressed a lower amount of *MyoD1* mRNA level compared to treated shSCRA myocytes (Supplementary Figure S3B). Since β-catenin was shown to induce *MyoD1* expression by binding the CER (43), chromatin immunoprecipitation (ChIP) was performed to evaluate the potential role of LSD1 in β-catenin recruitment on the *MyoD* CER during myoblast commitment. As expected, in shSCRA myoblast β-catenin was strongly enriched on the CER after 72 hours in MDM. Conversely shLSD1 myoblasts failed to recruit β-catenin on the CER (Figure 5B). Our data support the hypothesis that LSD1 is required for the recruitment of β-catenin on the *MyoD* CER.

**Figure 5.**
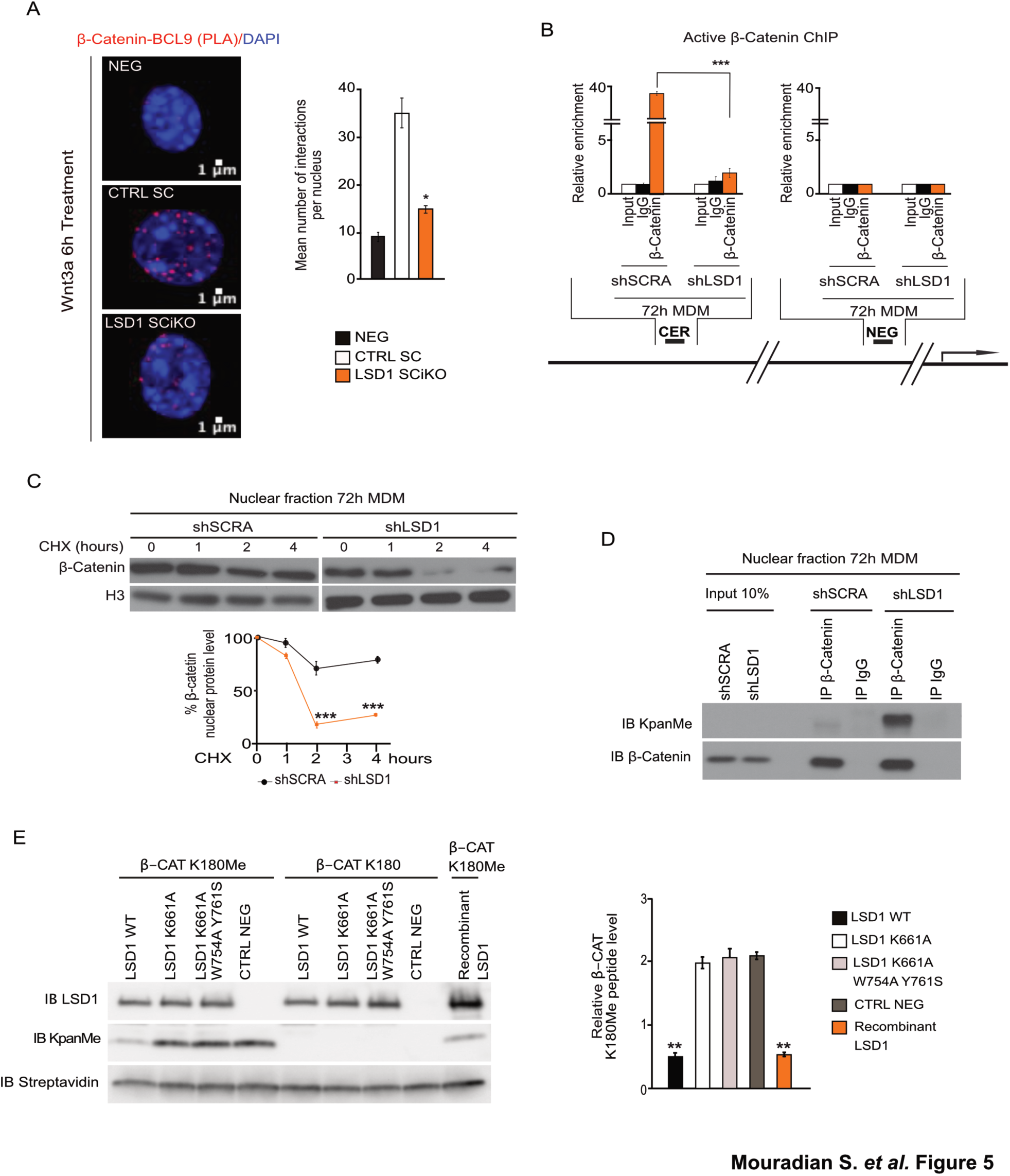
LSD1 demethylates β-catenin protein. (**A**) BCL-9 and β-catenin protein-protein interactions evaluated using *in situ* proximity ligation assay (PLA). Complexes visualized as red dots. Scale bar, 1 μm. Quantification of BCL-9/ β-catenin PLA assay on CTRL SC and LSD1 SCiKO cells after 6 h of Wnt3A treatment. Red dots were quantified in nucleus from at least 100 cells per condition. (**B**) Localization of β-catenin at the Core Enhancer region (CER) of *MyoD* gene locus after 72 h in myogenic differentiation medium (MDM). ChIP analysis was performed on shSCRA and shLSD1 cells with an anti-β-catenin antibody. Enrichment values were shown as fold difference relative to the NEG region. (**C**) LSD1 knockdown accelerated the turnover rate of endogenous β-catenin in shSCRA and shLSD1 cells after 72 h in MDM, in a time-course CHX treatment. (**D**) Loss of LSD1 function caused an increase in methylated β-catenin protein in the nucleus after 72h in MDM. (**E**) Demethylation assay using methylated and non-methylated lysine 180 β-catenin peptides as substrate. LSD1 WT, LSD1 K661A and LSD1 K661A/W754A/Y761S and commercial recombinant LSD1 were incubated with β-CAT K180Me and β-CAT K180 and analyzed by western blot with anti KpanMe antibody. The streptavidin antibody was used to detect the β-catenin peptides, which were conjugated to biotin. Values are mean of at least three experiments. ± SEM. *p < 0.05, **p < 0.01, ***p < 0.001 (Bonferroni test after one way-ANOVA).

### Demethylation by LSD1 is required for β-catenin stabilization in the nucleus

We next investigated the mechanism through which LSD1 promoted β-catenin activity. Whereas the levels of β-catenin transcript were similar in both cell lines (Supplementary Figure S3C), analysis of whole protein extracts by western blot showed that β-catenin levels were lower in shLSD1 than in shSCRA, (Supplementary Figure S3D). Of note, such reduction was due to the reduction of the levels of nuclear β-catenin, since cytosolic and membranous β-catenin levels were similar in the two conditions (Supplementary Figure S3E, F). A time-course experiment with cycloheximide (CHX) to evaluate the stability of nuclear β-catenin protein showed that β-catenin half-life was strongly reduced in shLSD1 cells compared to shSCRA cells (Figure 5C). Mono-methylation of β-catenin lysine 180 by SET7/9 destabilizes the protein (20). Consistently, upon treatment with the proteasome inhibitor MG132, non-methylable SET7/9 β-catenin mutant (β-CAT K180R) accumulates significantly less compared to the WT β-catenin (Supplementary Figure S3G).

Co-immunoprecipitation experiments indicated that both over-expressed and endogenous β-catenin interact with LSD1 (Supplementary Figure S3H, I). We thus speculated that LSD1 could directly demethylate β-catenin to stabilize it in the nucleus. β-catenin methylation was thus evaluated by western blot with an anti-pan methyl lysine (KpanMe) antibody on β-catenin immunoprecipitated from shSCRA or shLSD1 nuclear extracts. While in proliferating myoblasts nuclear β-catenin is methylated both in shLSD1 and shSCRA conditions (Supplementary Figure S3J), after 72h in MDM, nuclear β-catenin methylation was very low in shSCRA nuclei whereas it remained strongly methylated in shLSD1 nuclei, suggesting that β-catenin is demethylated by LSD1 during myoblast commitment (Figure 5D).

Finally, an *in vitro* de-methylation assay was performed with recombinant LSD1 to assess whether LSD1 directly demethylates β-catenin. A mono-methylated β-catenin peptide encompassing lysine 180 (β-CAT K180Me) was incubated with recombinant LSD1 or with cellular extracts of HEK 293T cells overexpressing either WT LSD1 or catalytically inactive LSD1 mutants (LSD1K661A and LSD1 K661A/W754A/Y761S, (44)). As shown in Figure 5E, both recombinant LSD1 and cellular extracts with WT LSD1 efficiently demethylated the β-cat K180Me peptide. Conversely, the catalytically inactive LSD1 mutants failed to demethylate the β-cat K180Me peptide. These results demonstrate that β-catenin is a non-histone substrate of LSD1 demethylase activity.

### LSD1 and β-catenin are involved in mitotic spindle orientation

Since β-catenin protein is required for the Wnt3A-dependent asymmetric cell division in ESCs (15), we checked whether this mechanism also applied to MuSCs. CTRL SC MuSCs were transiently transfected with an shRNA directed against β-catenin (shCTNNB1) and exposed to Wnt3A-coupled beads 24 h later. CTRL shCTNNB1 MuSCs exhibited mitotic spindle misorientation, with a significantly different angle distribution of the spindle compared to CTRL SC treated with Wnt3A-beads (Figure 4C and Figure 6A, C).

**Figure 6.**
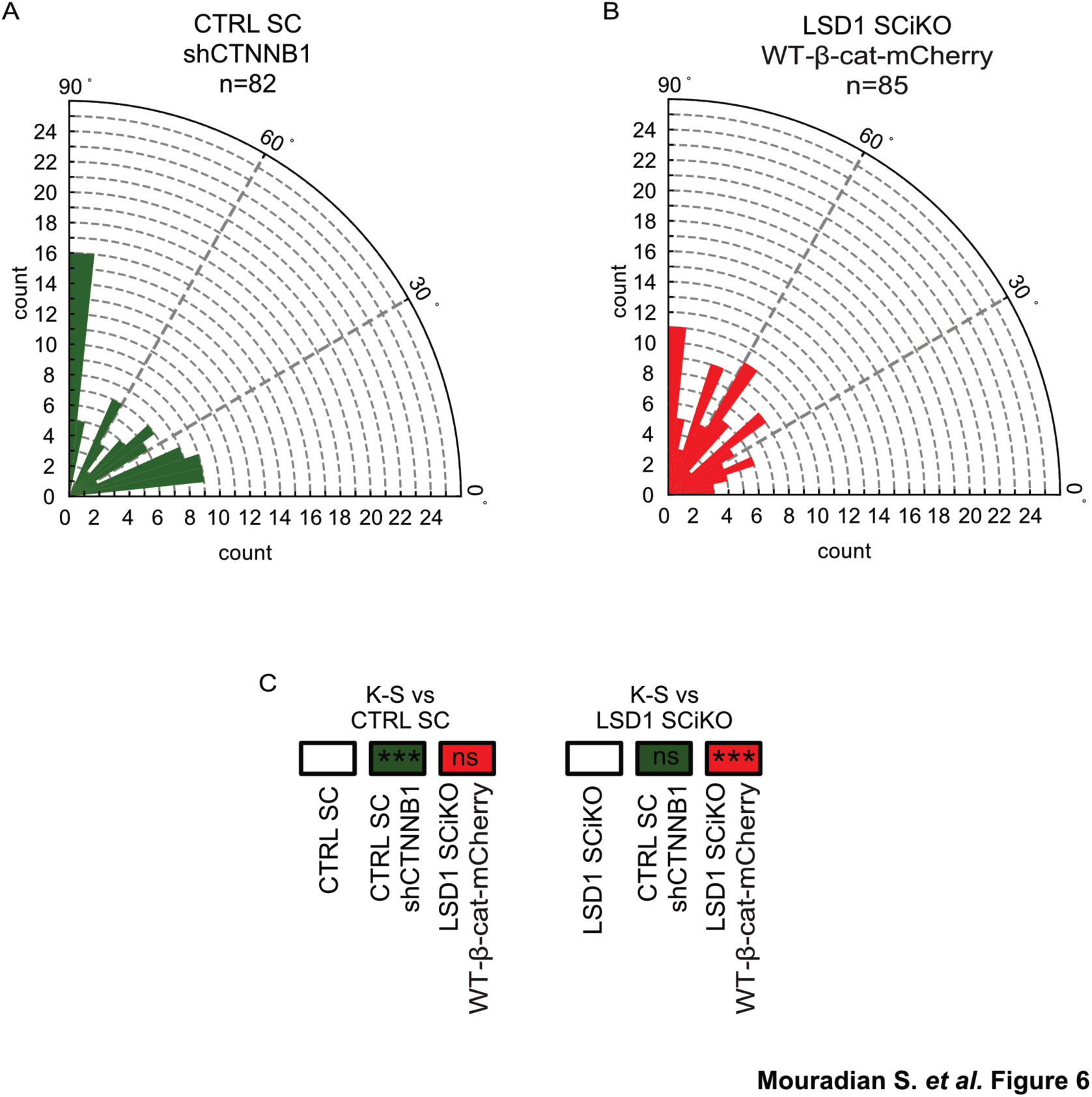
β-catenin is required for LSD1-mediated Wnt3A-spindle orientation. Rose plots depicting the distribution of mitotic spindle angle orientations in (**A**) CTRL SC treated with shRNA against *Ctnnb1* (shCTNNB1) or (**B**) LSD1 SCiKO MuSCs overexpressing *wild-type* β-catenin fused to mCherry (WT-β-cat-mCherry). n= number of cells. (**C**) ***p<0.001 in boxes indicate statistical significance calculated by multiple Kolmogorov-Smirnov tests against CTRL SC or LSD1 SCiKO MuSCs dividing with a Wnt3a-bead.

To further investigate if β-catenin over expression could compensate for the lack of LSD1, freshly isolated LSD1 SCiKO MuSCs were transfected with a plasmid encoding a *wild-type* β-catenin sequence fused with mCherry (WT-β-cat-mCherry). Indeed, β-catenin overexpression rescued the orientation of the mitotic spindles according to Wnt3A-beads in mCherry+ LSD1 SCiKO (Figure 6B, C). Altogether these results suggest that LSD1 involvement in mitotic spindle orientation is achieved through β-catenin.

### LSD1 controls β-catenin levels in stem cells

To investigate if the regulation of β-catenin stability by LSD1 also existed in non-muscle stem cells, LSD1 KO embryonic stem cell lines was generated by CRISPR/CAS9. CTRL WT and LSD1 KO ESCs displayed similar percentage of cells in S-Phase of the cell cycle (supplementary figure S4A). Upon Wnt3A treatment, while cytosolic β-catenin was similar between CTRL WT and LSD1 KO clones (supplementary figure S4B), we observed a severe decrease of nuclear β-catenin protein in LSD1 KO ESCs compared to CTRL WT ESCs (Figure 7A), accompanied by an impairment of β-catenin transcriptional activity (Figure 7B). Consistently with our results on MuSCs, a time-course experiment with CHX confirmed that in LSD1 KO ESCs nuclear β-catenin protein stability is significantly reduced compared to CTRL WT ESCs (Figure 7C). We then exposed these ESCs to Wnt3A-coupled beads and we measured the orientation of the mitotic spindle towards the Wnt3A-coupled beads. Consistent with previous report (14, 15), CTRL WT ESCs oriented toward the Wnt3A-coupled beads with an angle of 85-90° (Figure 7D). Conversely the majority of LSD1 KO ESCs divided with an angle lower than 30° (Figure 7E, F), phenocopying the β-catenin KO mitotic spindle misorientation (Supplementary figure S4C, D) (15). Altogether, these results indicate that the regulation of β-catenin by LSD1 is not cell type-specific.

**Figure 7.**
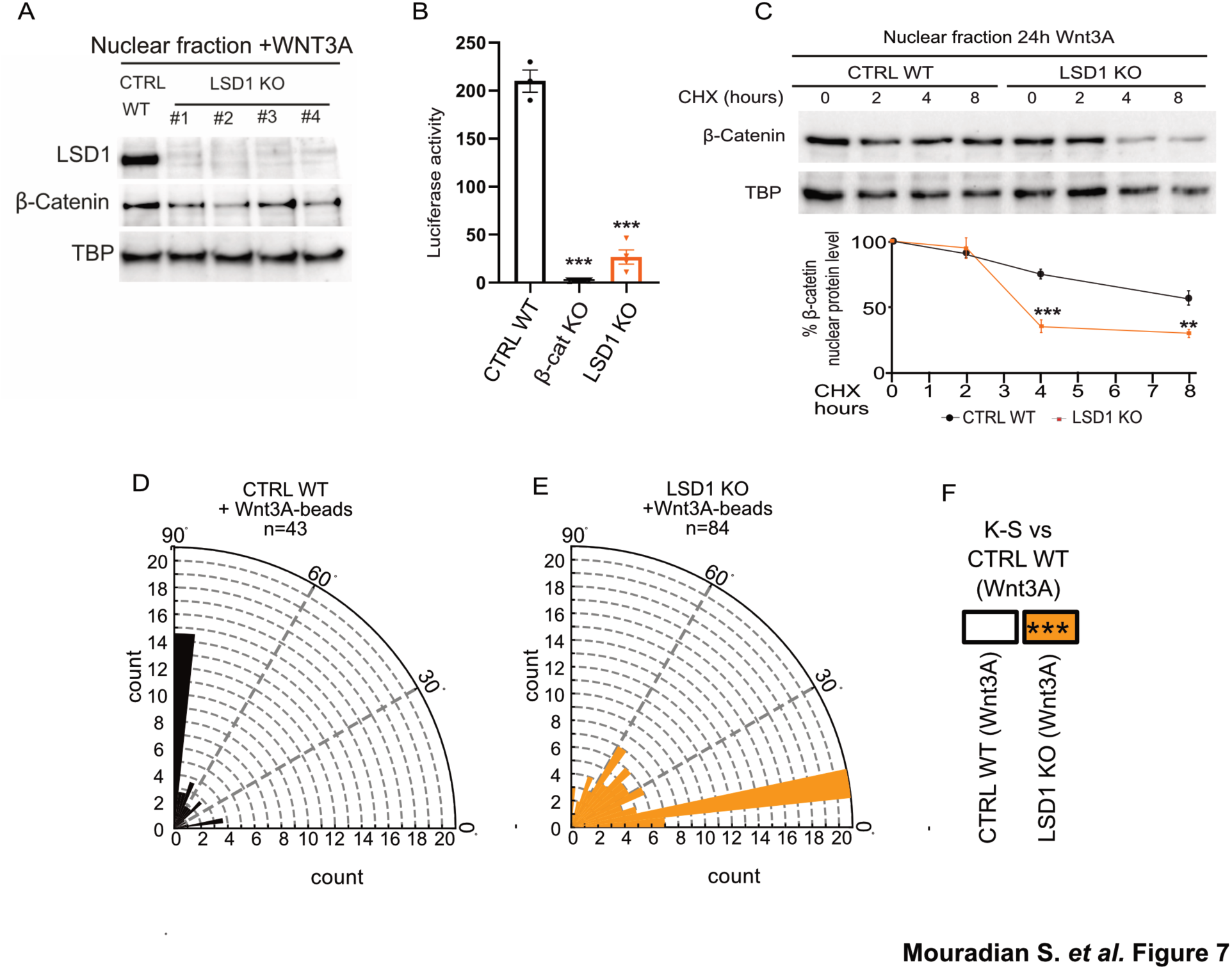
LSD1 regulates β-catenin turnover in ESCs. (**A**) Western blot analysis of nuclear β-catenin protein in CTRL WT and 4 different LSD1 KO ESCs clones. (**B**) The TCF transcriptional activity of CTRL WT, β-Cat KO and LSD1 KO ESCs is shown as a ratio of TOP-FLASH to FOP-FLASH luciferase-mediated signals, when cultured for 6 h in the presence of Wnt3A. (**C**) Upon Wnt3A treatment, LSD1 KO ESCs displayed an accelerated turnover rate of nuclear β-catenin compared to CTRL WT ESC, in a time-course CHX treatment. Values are mean of at least three experiments. ± SEM. **p < 0.01, ***p < 0.001 (Bonferroni test after one way-ANOVA). Rose plots depicting the distribution of mitotic spindle angle orientations in (**D**) CTRL WT or (**E**) LSD1 KO ESCs. n= number of cells. (**F**) ***p<0.001 in box indicates statistical significance calculated by multiple Kolmogorov-Smirnov tests against CTRL WT ESC dividing with a Wnt3a-beads.

## Discussion

Although canonical Wnt/β-Catenin signaling pathway has long been known to play a key role in MuSCs fate during skeletal muscle regeneration, its mechanism of action is not fully characterized. In this study, we demonstrate that LSD1 is required to stabilize nuclear β-catenin by demethylating the lysine 180. This enables β-catenin to be recruited on the *MyoD1* Core enhancer region and initiate the MuSCs commitment to differentiation. We also provide evidence that LSD1 favors Wnt3A-mediated mitotic spindle orientation in MuSCs and ESCs via β-catenin. So far, LSD1 is the first demethylase identified to directly target β-catenin protein and influence its activity.

The first methyltransferase identified to mono-methylate β-catenin on K180 is SET7/9 (19). β-catenin K180me1 destabilizes the protein, which is degraded faster (20). Another study conducted on MuSCs showed that SET7/9 activity is required for β-catenin localization in the nucleus rather than for its stability (17). However, the authors did not demonstrate a direct methyltransferase activity of SET7/9 on β-catenin protein (17). These results raised the possibility that the effect of SET7/9 methylation on β-catenin might be cell type– and cell compartment-specific. Our study shows that depletion of LSD1 causes a reduction of nuclear β-catenin but has no effect on cytoplasmic or membranous β-catenin during MuSCs commitment. Thus, the role of LSD1 is to prevent β-catenin degradation once it has entered in the nucleus. Our results also show that in embryonic stem cells, LSD1 is required to maintain β-catenin level and transcriptional activity, indicating that the action of LSD1 on β-catenin is not restricted to muscle stem cells.

It has been recently demonstrated that in myoblast cultures, LSD1 inhibition could promote the switch of myogenic cells into brown adipocytes in pro-adipogenic conditions (45). Our results indicate that in pro-myogenic conditions, MuSCs, lacking LSD1, do not lose their myogenic identity. Similarly, we show that LSD1 inactivation *in vivo* did not induced adipogenesis. Both genetic inactivation and the pharmacological inhibition of LSD1 led to a significative expansion of PAX7+ MuSCs after muscle injury, which remained fully myogenic even after repeated injuries.

To maintain MuSCs homeostasis and generate appropriate numbers of transient amplifying progenitors to support the regeneration of muscle, MuSCs can undergo asymmetric (ACD) or proliferating/self-renewing symmetric (PSCD) or differentiation symmetric (DSCD) division. LSD1 inactivation in MuSCs increases the probability to undergo proliferating/self-renewing symmetric division resulting in an expansion of progenitor cells able to repair muscle fiber and to self-renew. MuSCs ability to switch between asymmetric or symmetric divisions is linked to the orientation of the mitotic spindle and is finely controlled by extrinsic signaling pathways that regulate intrinsic fate determinants. Taking advantage of a method published recently (40), we demonstrate that Wnt/β-catenin signaling pathway and LSD1 are implicated in MuSCs ACD choice over PSCD *in vitro*. Indeed, either β-catenin knock down or LSD1 inactivation in MuSCs result in the loss of mitotic spindle orientation toward the Wnt3A-beads. Moreover, β-catenin overexpression in LSD1 KO MuSCs restores mitotic spindle orientation by Wnt3A. These results place LSD1 as a crucial partner of β-catenin for Wnt3A-mediated mitotic spindle orientation in muscle stem cells.

β-catenin loss of function studies have shown that β-catenin is required to maintain the regenerative potential of MuSCs (22). In our LSD1 SCiKO mouse model, despite an early delay in MuSCs commitment, the regeneration process finally occurs. These different phenotypes could be explained by the complete absence of β-catenin protein (at the membrane, in the cytosol and in the nucleus) in the β-catenin KO model (22), while in the LSD1 SCiKO model, β-catenin levels are specifically reduced only in the nucleus. This suggests that β-catenin might be involved in non-nuclear function independently from LSD1 during skeletal muscle regeneration. We therefore propose a regulation of nuclear β-catenin turnover by LSD1 to be necessary for the transcription of the *MyoD1* CER. In agreement with our model, β-catenin KO MuSCs as well as SET7/9 KO MuSCs show a significantly lower expression of *MyoD1* gene (17, 22). It would be interesting to compare the Wnt3A-mediated spindle orientation distribution between LSD1 SCiKO and these two others models, supported by our evidence that β-catenin silencing in MuSCs drastically mis-oriented the mitotic spindle distribution.

Interestingly, we demonstrate that LSD1/β-catenin axis is not a MuSCs-specific molecular mechanism. Indeed, knocking out LSD1 in ESCs causes a reduction of β-catenin nuclear protein level and a mitotic spindle mis-orientation upon Wnt3A treatment, mimicking the phenotype observed in β-catenin KO ESC (15). Upon Wnt3A treatment, β-catenin KO ESCs show an aberrant symmetric segregation of pluripotency markers, leading to an impairment of ESC differentiation. Remarkably, whole genome distribution studies have shown the same phenotype in LSD1 KO ESCs, where absence of LSD1 activity leads to a re-expression of pluripotent genes in differentiating ESCs (46). Collectively these findings support the hypothesis that the regulation of enhancers by LSD1 involves a dual mechanism: modification of histones and stabilization of nuclear β-catenin.

In conclusion, our results identify LSD1 as a new checkpoint in the canonical Wnt/β-catenin pathway, required for transcriptional regulation by β-catenin and Wnt3A-mediated mitotic spindle orientation. It will be interesting to investigate the correlation between LSD1 and β-catenin in diseases, such as colorectal cancer, characterized by both LSD1 overexpression and an aberrant activation and hence accumulation of nuclear β-catenin. The findings might elucidate important mechanisms underlying the development of these diseases from a different point of view.

## Data availability

All data generated or analyzed during this study are included in this published article and its supplementary file.

## Acknowledgments

We thank Céline Angleraux and Pierre Contard from the PBES in the SFR biosciences (UMS3444) for animal breeding. We thank T. Andrieu and S. Dussurgey from the AniRA-Cytometry platform in the SFR biosciences (UMS3444) for their expertise in cell sorting, and D. Ressnikoff and B. Chapuis (CIQLE imaging center, SFR Santé Lyon-Est (UMS3453) for their help with image acquisition. We thank Dr. Denise Aubert for kindly provided ESCs *wild-type*. We are also grateful with Dr. Marta Shahbazi and Dr. Shukry Habib for having shared the heterozygous β-catenin deficient (βfl/-) ESCs. We also thank R. Mounier for critical reading the manuscript.

## Author Contributions

I.S. and L.S. conceived the research. S.M. and D.C. made *Lsd1* muscle specific inactivation. S.M. performed the CTX muscle injury. S.M., D.C. and F.B. performed immunofluorescence and histology on cryosections. V.R. generated LSD1 KO ESC model. F.L.G and N.R. contribute to Figure 1B. T.S. performed Rose plot visualization. N.L. and I.S. performed the biochemical experiments and the revisions. I.S. performed and analyzed myofiber culture and Wnt3a-dependent asymmetric division. I.S. wrote the manuscript with comments from all the authors.

