## Supplementary data for "LSD1 controls a nuclear checkpoint in Wnt/β-Catenin signaling to regulate muscle stem cell self-renewal"

This file contains following material:

**Supplemental Table S1.** DNA/RNA oligonucleotides, antibodies, plasmids and mouse models used in this study.
**Figure S1.** LSD1 inactivation delays myogenic differentiation.
**Figure S2.** LSD1 inactivation affects early step of muscle regeneration.
**Figure S3.** LSD1 is required for β-catenin transcriptional activity.
**Figure S4.** LSD1 inactivation does not influence ESC maintenance.

**Supplemental Table S1.** DNA/RNA oligonucleotides, plasmids, antibodies and mouse models used in this study.

| REAGENT or RESOURCE | SOURCE | IDENTIFIER |
|---|---|---|
| **<u>Experimental models: organisms/strains:</u>** | | |
| C57BL/6J mice | Charles River | N/A |
| Pax7<tm2.1(cre/ERT2)Fan>/J | The Jackson Laboratories | #012476 |
| LSD1tm1Schüle | Zhu, D et al., Nat commun 2014 | N/A |
| **<u>Oligonucleotides:</u>** | | |
| LSD1 Wt For genotyping: ATA-CGA-AGT-TAT-GGA-TCC-AAG | Eurogentec | N/A |
| LSD1 Flox For genotyping: CCT-ACA-CTG-TGC-CAG-GCT-GC | Eurogentec | N/A |
| LSD1 Rev genotyping: GCA-GGC-GGT-TTG-AAA-TGT-ATT-C | Eurogentec | N/A |
| Cre For genotyping: CGA-TGC-AAC-GAG-TGA-TGA-GG | Eurogentec | N/A |
| Cre Rev genotyping : GCA-TTG-CTG-TCA-CTT-GGT-CGT | Eurogentec | N/A |
| *MyoD* NEG For ChIP: CCC-TTC-ATC-CAG-GGC-ACT-AC | Eurogentec | N/A |
| *MyoD* NEG Rev ChIP: TTG-GGA-ACC-CAG-CAG-TAA-GC | Eurogentec | N/A |
| *MyoD* CER For ChIP: CTA-AAC-ACC-AGG-CAT-GAG-AGG | Eurogentec | N/A |
| *MyoD* CER Rev ChIP: ACT-CAC-TTT-CTC-CCA-GAG-TTG-C | Eurogentec | N/A |
| *Fst* For Real-Time qPCR: CTG-CTG-CTA-CTC-TGC-CAG-TT | Eurogentec | N/A |
| *Fst* Rev Real-Time qPCR: ACA-TCC-TCC-TCG-GTC-CA-TGA | Eurogentec | N/A |
| *Axin2* For Real-Time qPCR: GGG-TTC-TGA-AAT-TCA-TAG-ACT | Eurogentec | N/A |
| *Axin2* Rev Real-Time qPCR: CGA-CTG-TTC-AAT-AAA-TAT-CAG | Eurogentec | N/A |
| *Ctnnb1* For Real-Time qPCR: TAC-GAG-CAC-ATC-AGG-ACA-CC | Eurogentec | N/A |
| *Ctnnb1* Rev Real-Time qPCR: ACA-ATC-CGG-TTG-TGA-ACG-TC | Eurogentec | N/A |
| *MyoD1* For Real-Time qPCR: AGC-ACT-ACA-GTG-GCG-ACT-CA | Eurogentec | N/A |

| | | |
|---|---|---|
| *MyoD1* Rev Real-Time qPCR: GCT-CCA-CTA-TGC-TGG-ACA-GG | Eurogentec | N/A |
| **Plasmids:** | | |
| pCMX-LSD1 flag | Laboratory of Roland Schüle | N/A |
| pCMX-LSD1 K661A flag | Laboratory of Roland Schüle | N/A |
| pCMX-LSD1 K661A/W754A/Y761A flag | Laboratory of Roland Schüle | N/A |
| pCMV-GFP | GenScript | N/A |
| pCMV β-CAT | GenScript | N/A |
| pCMV β-CAT K180R | GenScript | N/A |
| M50 super 8xTOPFLASH | Addgene | #12456 |
| M51 super 8xFOPFLASH | Addgene | #12457 |
| pRL-TK | Promega | #E2241 |
| shRNA against CTNNB1 | Merck | TRCN0000012690 |
| CTNNB1 WT_mCherry_pCAGIG | GenScript | N/A |
| **Recombinant proteins:** | | |
| β-CAT WT: Biotin-GGGGGAAVMVHQLSKKEASRHAIMRSPQMVSAIVRTMQNTNDVETARCTAGTLHNLSHHREGLLAIF | Proteogenix SAS | N/A |
| β-CAT K180me: Biotin-GGGGGAAVMVHQLSK(me)KEASRHAIMRSPQMV SAIVRTMQNTNDVETARCTAGTLHNLSHHREGLLAIF | Proteogenix SAS | N/A |
| Flag: MDYKDHDGDYKDHDIDYKDDDDK | Proteogenix SAS | N/A |
| Recombinant LSD1 | Merck | #SRP0122 |
| **Antibodies:** | | |
| Mouse anti-Pax7 | DSHB | #Pax7 |
| Rabbit anti-Ki67 | Cell Signaling | #9129 |
| Rabbit anti-LSD1 | Abcam | #17721 |
| Rabbit anti-Laminin | Merck | #L9393 |
| Mouse anti-MYF5 | Active motif | # 39801 |
| Mouse anti-Myogenin | DSHB | #F5D |
| Rabbit anti-pan methyl Lysine | Abcam | #7315 |
| Rabbit anti-GFP | Merck | #G1544 |
| Streptavidin-HRP | Thermo Fisher scientific | #434323 |
| Monoclonal ANTI-FLAG® M2 | Merck | #F1804 |

| | | |
|---|---|---|
| Rabbit anti- Non-phospho (Active) β-catenin | Cell Signaling | #8814 |
| Rabbit anti-β-catenin | Cell Signaling | #9562S |
| Rabbit anti-H3 | Cell Signaling | #4499S |
| Rabbit anti-BCL9 | Invitrogen | #PA5-49466 |
| Rabbit anti-Desmin | Cell Signaling | #5332 |
| Mouse anti-Myosin | DSHB | #A4.1025 |
| Rat anti-CD34-FITC | Thermo Fisher scientific | #11-0341-82 |
| Rat anti-CD45-PE | Thermo Fisher scientific | #12-0451-82 |
| Rat anti-CD31-PE | Thermo Fisher scientific | #12-0311-82 |
| Rat anti-Ly-6A/E(Sca-1) -PE | Thermo Fisher scientific | #12-5981-82 |
| Rat anti-Alpha 7 integrin 647 | AbLab | #67-0010-05 |
| Alexa Fluor 488-conjugated Goat Anti mouse IgG | Jackson lab | #115-486-072 |
| Alexa Fluor 546-conjugated Donkey Anti Rabbit IgG | Molecular probes | #A10040 |

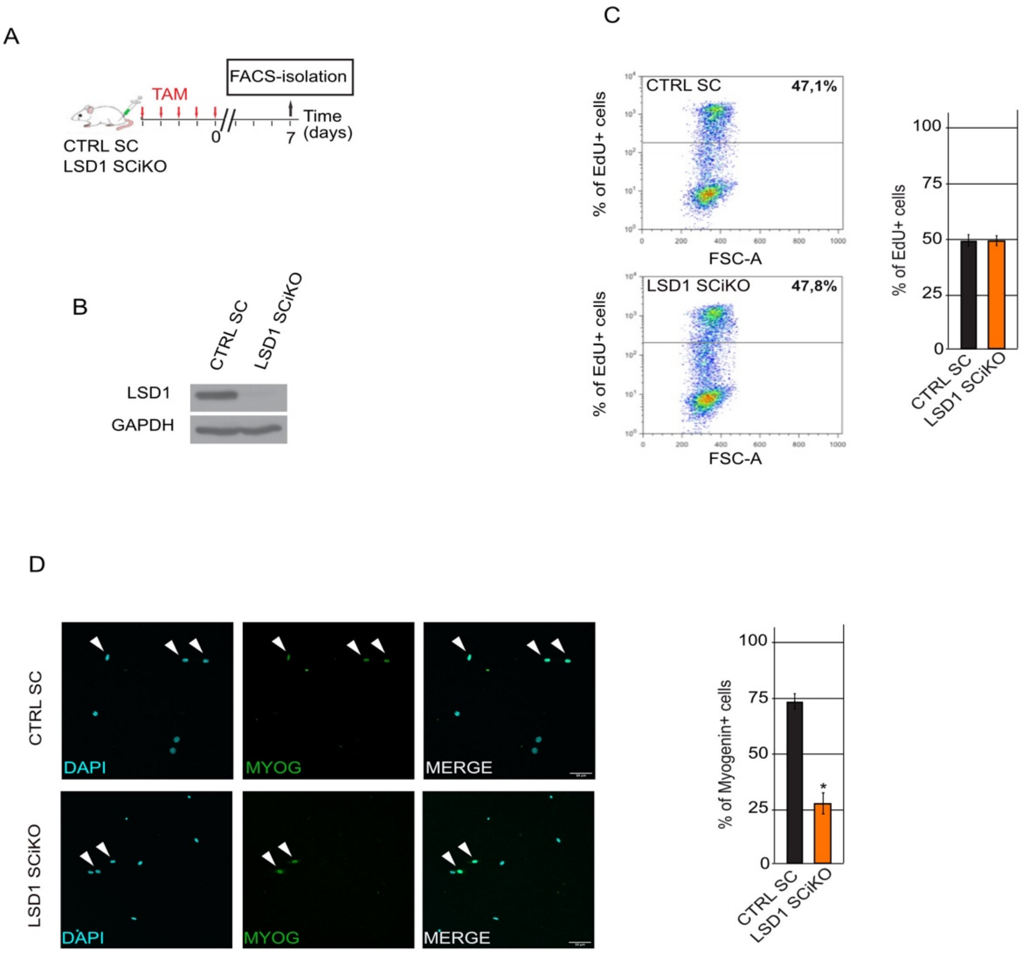


**Figure S1. LSD1 inactivation delays myogenic differentiation *in vitro*.** (**A**) Experimental set up. (**B**) LSD1 immunoblot on FACS-sorted MuSCs from CTRL SC and LSD1 SCiKO muscles cultured in growth medium for 4 days. GAPDH was used as loading control. (**C**) Percentage of CTRL SC and LSD1 SCiKO MuSCs in S-Phase. Measurements were made by cytometry analysis after treatment with EdU for two hours. (**D**) Myogenin immunostaining of FACS-sorted MuSCs from CTRL SC and LSD1 SCiKO muscles, seeded at low density, after 48 h of myogenic differentiation medium. Representative images of MYOG positive MuSCs are shown.
Scale bars, 50 µm. n = 5 primary cell cultures/genotype. Values are percentage mean ± SEM. *$p < 0.05$ (Bonferroni test after one way-ANOVA).

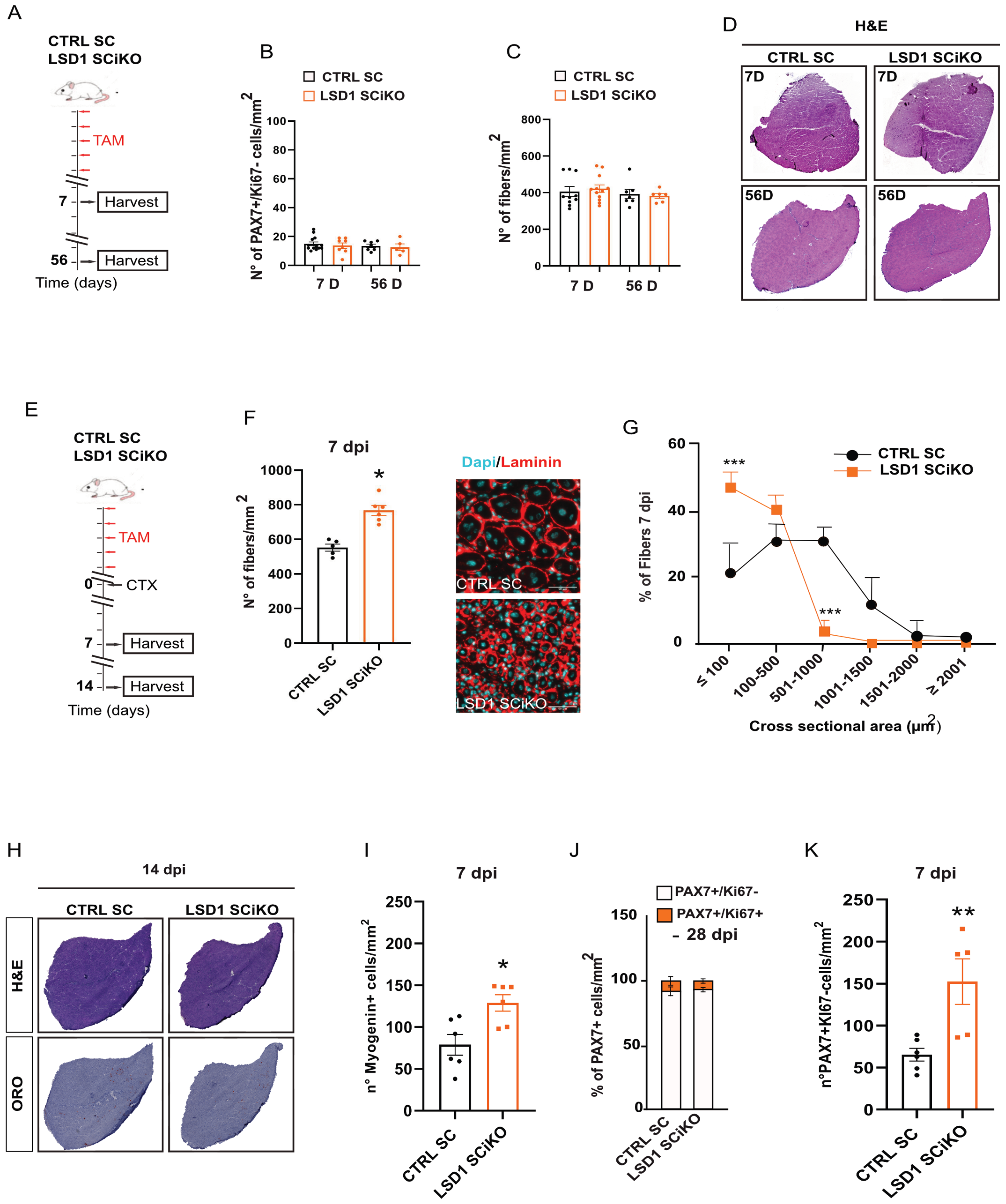


**Figure S2. LSD1 inactivation affects early step of muscle regeneration.** (**A**) Experimental set up. (**B**) Quantification of the number of sublaminar PAX7+/Ki67- cells per mm$^2$. (**C**) Quantification of the number of myofibers per mm$^2$. (**D**) Histological analysis of TA muscle in CTRL SC and LSD1 SCiKO mice. (**E**) CTX experimental setup. (**F**) Anti-Laminin staining on cryosections of regenerated TA muscles in CTRL SC and LSD1 SCiKO mice at 7 dpi. Quantification of the number of myofibers per mm$^2$. (**G**) CSA distribution of muscle fibers in CTRL SC and LSD1 SCiKO mice TA cryosections at 7 dpi. (**H**) Hematoxylin-eosin and Oil Red O staining on cryosections of regenerated TA muscles in CTRL SC and LSD1 SCiKO mice at 14 dpi. (**I**) Quantification of the number of Myogenin + cells per mm$^2$ in CTRL SC and LSD1 SCiKO mice at 7 dpi. (**J**) Percentage of PAX7+/Ki67+ (orange, proliferating cells)

MuSCs at 28 dpi was quantified per $mm^2$. (**K**) Quantification of the number of PAX7+/Ki67- cells per $mm^2$ in CTRL SC and LSD1 SCiKO mice at 7 dpi.
Scale bars, 50 µm. n = 4 mice/genotype. Values are mean or percentage mean ± SEM. *p < 0.05, **p < 0.006, ***p < 0.001 (Bonferroni test after one way-ANOVA).

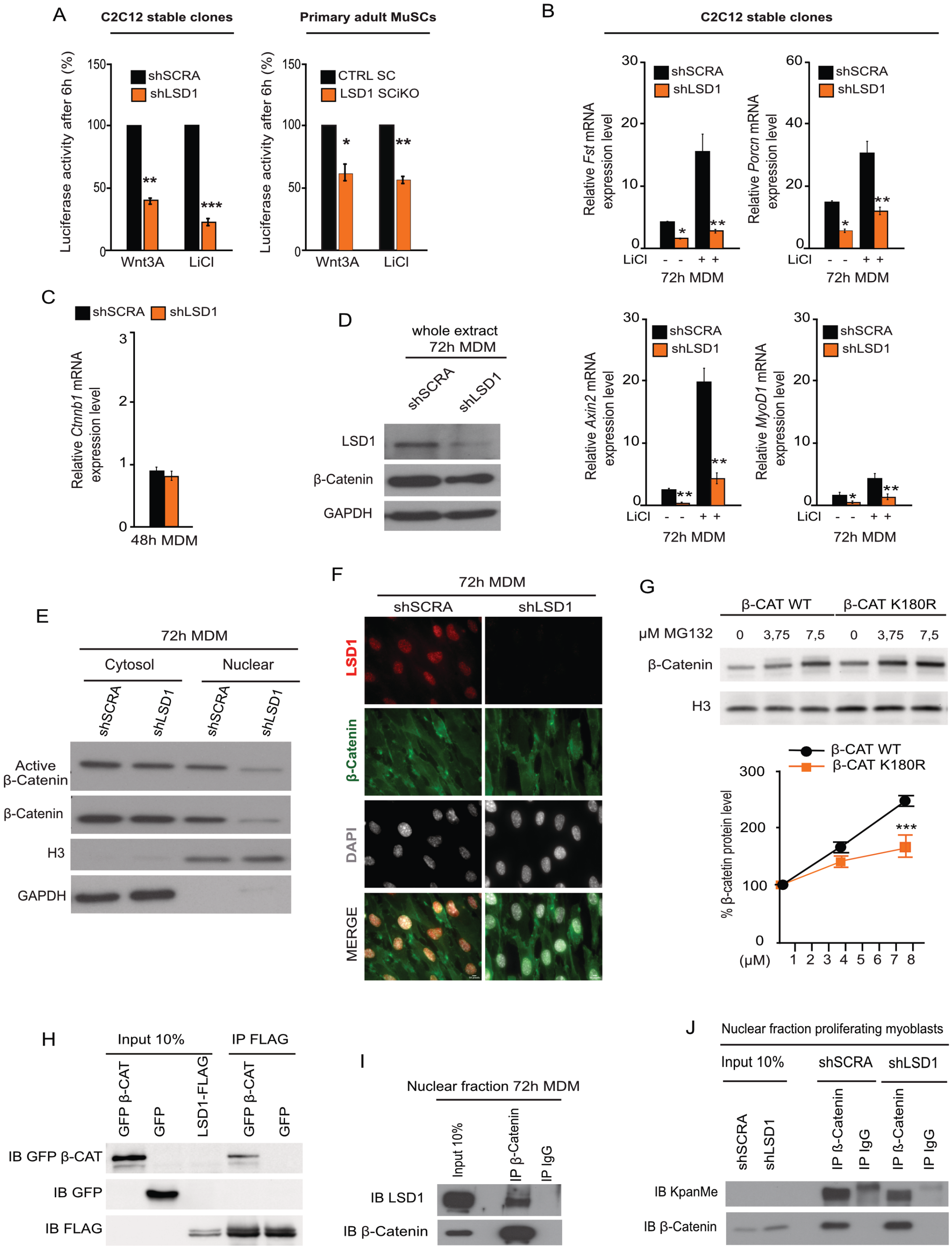


**Figure S3. LSD1 is required for β-catenin transcriptional activity.** (**A**) The TCF transcriptional activity of C2C12 (shSCRA and shLSD1) and MuSC (CTRL SC and LSD1 SCiKO) cells is shown as a ratio of TOP-FLASH to FOP-FLASH luciferase-mediated signals, when cultured for 6 h in the presence of Wnt3A or LiCl. (**B**) *Fst, Porcn, Axin2* and *MyoD1* mRNA levels in shSCRA and shLSD1 cells untreated and treated with LiCl during differentiation. RT-qPCR values were normalized to the *Ppib* mRNA. mRNA levels are shown as the fold variation compared to shSCRA cells at MDM0. (**C**) *Ctnnb1* mRNA levels in shSCRA and shLSD1 cells after 48 h in MDM. RT-qPCR values were normalized to the *Ppib* mRNA. mRNA levels are shown as the fold

variation compared to shSCRA cells at MDM0. (**D**) Western blot analysis of β-catenin in whole protein extract of shSCRA and shLSD1 cells after 72 h in MDM. GAPDH was used as loading control. (**E**) Western-blot analysis of β-catenin in the two cellular compartments (Cytosol: GAPDH) and Nucleus (H3). (**F**) Anti-LSD1 and anti- β-catenin immunostaining of shSCRA and shLSD1 cells after 72h in MDM. (**G**) Detection of β-catenin WT or mutant β-catenin K180R protein level in HEK 393T cells without or with treatment with MG132. (**H**) CoIP detection of the binding of LSD1 to β-catenin in transfected HEK 293T cells. (**I**) Endogenous β-catenin and LSD1 interacted with each other in the nuclear fraction of C2C12 cells after 72 h in MDM. (**J**) LSD1 does not affect the β-catenin protein methylation status in proliferating myoblasts.
Values are mean or percentage mean of at least three experiments. ± SEM. *$p < 0.05$, **$p < 0.01$, ***$p < 0.001$ (Bonferroni test after one way-ANOVA).

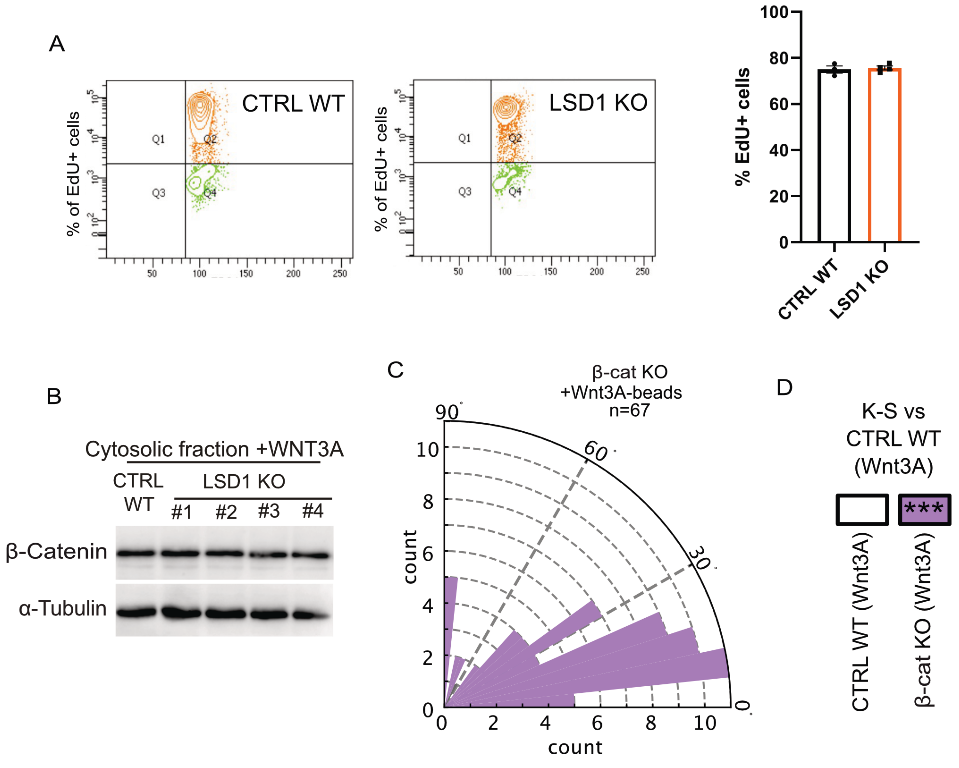


**Figure S4. LSD1 inactivation does not influence ESC maintenance.** (**A**) Percentage of CTRL WT and LSD1 KO ESCs in S-Phase. Measurements were made by cytometry analysis after treatment with EdU for two hours. (**B**) Western blot analysis of cytosolic β-catenin protein in CTRL WT and 4 different LSD1 KO ESCs clones. (**C**) Rose plot depicting the distribution of mitotic spindle angle orientations in β-catenin KO ESCs. n= number of cells. (**D**) ***p<0.001 in box indicates statistical significance calculated by multiple Kolmogorov-Smirnov tests against CTRL WT ESC dividing with a Wnt3a-bead.
